# Optimization of chondroitin production in *E. coli* using genome scale models

**DOI:** 10.1101/2023.12.22.573033

**Authors:** Márcia R. Couto, Joana L. Rodrigues, Adelaide Braga, Oscar Dias, Lígia Rodrigues

## Abstract

Chondroitin is a natural occurring glycosaminoglycan with applications as a nutraceutical and pharmaceutical ingredient and can be extracted from animal tissues. Microbial chondroitin-like polysaccharides emerged as a safer and more sustainable alternative source. However, chondroitin titers using either natural or recombinant microorganisms are still far from meeting the increasing demand. The use of genome-scale models and computational predictions can assist the design of microbial cell factories with possible improved titers of these value-added compounds. Genome-scale models have been used to predict genetic modifications in *Escherichia coli* engineered strains that would potentially lead to improved chondroitin production. Additionally, using synthetic biology approaches, a pathway for producing chondroitin has been designed and engineered in *E. coli*. Afterwards, the most promising mutants identified based on bioinformatics predictions were constructed and evaluated for chondroitin production in flask fermentation. This resulted in the production of 118 mg/L of extracellular chondroitin by overexpressing both superoxide dismutase (*sodA*) and a lytic murein transglycosylase (*mltB*). Then, batch and fed-batch fermentations at bioreactor scale were also evaluated, in which the mutant overexpressing *mltB* led to an extracellular chondroitin production of 427 mg/L and 535 mg/L, respectively. The computational approach herein described identified several potential novel targets for improved chondroitin biosynthesis, which may ultimately lead to a more efficient production of this glycosaminoglycan.

## Introduction

Genome-scale metabolic models (GEMs) are mathematical representations of the entire metabolic network of an organism (or consortium), that include a description of genes, enzymes, reactions, metabolites, as well as their associations and compartments, ultimately allowing to predict biological capabilities ^1^. Along with the increasing knowledge and data provided by genome sequencing technologies, the number, quality, and applications of available GEMs have been growing. These models provide valuable information for metabolic engineering strategies as they allow to predict phenotypic behavior of either wild-type or mutated strains under different environmental conditions and to identify targets to improve the metabolic flux towards a target product ^2,3^. Applications of GEMs on drug target identification/ drug discovery/ drug design and human disease studying have also been described ^2^. The BiGG Models repository ^4^ currently provides 108 open-source manually curated GEMs, that exhibit robustness of growth predictions.

GEMs representation follows a matrix configuration where the columns represent reactions and the rows metabolites (stoichiometric matrix). For each metabolite, its stoichiometric coefficient in each reaction is presented. Negative coefficients correspond to consumed substrates (reactants), and positive coefficients represent produced metabolites (products). The bounds for each reaction are defined to constrain the range of possible fluxes. The phenotype can then be estimated by different tools. Flux balance analysis (FBA) is a common mathematical method for analyzing the flux distributions through the metabolic network. This approach relies on mass balance equations assuming steady-state growth (all mass that enters the system must leave, so no metabolite is accumulated)^1^. Considering the constraints imposed by mass balance equations, environmental conditions and model metabolic bounds, FBA computes a possible flux distribution by maximizing an objective function by linear programming. Commonly, the biomass production reaction/equation is used as the objective function for simulating microbial phenotypes. FBA accurately predicts wild-type behavior, although for mutant phenotype prediction the simulation method Minimization of Metabolic Adjustment (MOMA) ^5^ and linear MOMA (LMOMA) ^6,7^ have been more frequently used. These methods assume that engineered strains will no longer grow to optimize biomass, but instead they grow to maintain the flux distribution as close to the wild-type as possible.

While simulation methods are essential for phenotype prediction, finding combinations of genetic modifications to reach a desired phenotype requires more complex computational tools to iteratively generate and evaluate candidate solutions until a desired phenotype or other termination criteria is achieved ^8^. Examples of such computational strain optimization methods include OptKnock ^9^, OptStrain ^10^, OptGene ^11^, simulated annealing/ evolutionary algorithms (SA/SEA) ^12^, FluxDesign ^13^, OptORF ^14^ and OptForce ^15^. These tools differ in the defined objective function, optimization algorithms, simulation methods, the information they support, and/or in the mathematical formulation ^8^. Biological behavior prediction through computational modulation has been widely and successfully used to increase the biotechnological production of several high-value compounds such as fatty acids ^16^, organic acids ^17–20^, lipids ^21–23^, polymers ^3,24–26^, amino acids ^27^, butanol ^28^, naringenin ^29^ and the glycosaminoglycan hyaluronic acid ^30^.

As reviewed in ^31^, chondroitin is a glycosaminoglycan (GAG) with several applications mainly used as a chondroprotective ingredient in human and veterinary medical prescriptions ^31^. The usual chondroitin source in current chondroitin-based market products is cartilages such as shark or cows, and efforts are being implemented to shift the production process to a more sustainable and safer one such as the biological production using microorganisms ^31^. The biotechnological production of chondroitin is still challenging mainly due to the low titers obtained ^32^. Strategies to improve microbial strains for producing chondroitin are therefore essential. *E. coli* has been the most widely used and well-known host for biotechnological applications, participating as a living catalyst in well-established industrial processes. Therefore, this microorganism was herein used as the host for heterologous chondroitin production and *in silico* strategies have been developed to find potential gene manipulation targets.

## Experimental

### Model construction and computational approach

*E. coli* BL21’s stoichiometric models iEC1356_Bl21DE3, iB21_1397, iECBD_1354 and the parent model iJO1366, from *E. coli* K-12, were obtained from the BiGG Models database (http://bigg.ucsd.edu/). These models were modified to include the heterologous pathway for chondroitin production *in silico*. The included reactions were uridine-diphosphate (UDP)-*N*-acetylglucosamine 4-epimerase (UAE), chondroitin synthase/polymerase (CHSY) and their corresponding genes, as well as a chondroitin exchange reaction. Chondroitin was added as new species in all models, and UDP-*N*-acetyl-D-galactosamine was also included, except for iEC1356_Bl21DE3 model that already contained this intermediate. The resulting models harboring the chondroitin pathway were named iEC1356_Bl21DE3_c, iB21_1397_c, iECBD_1354_c and iJO1366_c.

The modified models were uploaded/imported in the workbench OptFlux Version 3.3.5 ^33^. The Linear Programming solver employed in this study was CPLEX Optimization Studio Version 12.9.0, developed by IBM. Evolutionary optimization was performed for gene deletion and for gene under and overexpression predictions, to search for mutants with enhanced flux for the chondroitin production reaction. The optimization method employed in this study was the Strength Pareto Evolutionary Algorithm 2 (SPEA2). Optimizations were performed with parsimonious FBA (pFBA) ^34^ and using the Biomass-Product Coupled Yield (BPCY) as the objective function. The maximum for evaluation functions was set to 50000. A maximum of 10 modifications was allowed. The environmental conditions were set to aerobic (without any restriction of oxygen, lower bound set to -1000) and the substrate glucose to 10 mmol/gDW/h (lower bound set to -10). A recently described alternative computational approach was also performed using the MEWpy workbench developed for Python ^35^. *E. coli* BL21 model iB21_1397 was used for this approach. Under the identical environmental conditions as previously described, the evolutionary optimization was conducted using a multi-objective approach. The objectives were set as the BPCY and the Weighted Yield (WYIELD), representing the weighed sum of the minimum and maximum product fluxes. pFBA was used as the phenotype prediction method and the evolutionary algorithm (EA) Non-dominated Sorting Genetic Algorithm II ^36^ was used for the optimization approach.

To evaluate the robustness of the obtained solutions from strain optimization, flux variability analysis (FVA) was performed by fixing biomass as the obtained in the mutant solution and alternating the pFBA objective function between maximizing and minimizing chondroitin production ^37^.

### Heterologous pathway construction

The strains and plasmids used to construct a chondroitin production pathway *in vivo* in *E. coli* are described in Table 1. Two reactions were included in *E. coli* enzymatic machinery, namely UAE and CHSY. The selected genes to perform these enzymatic reactions were the ones from the pathogenic strain *E. coli* O5:K4:H4 which naturally produces a compound analogue to chondroitin as part of its capsule ^38^. The genes *kfoA* and *kfoC* (encoding UAE and CHSY, respectively) present in the pETM6 plasmid, were kindly provided by Dr. Matheos Koffas (Rensselaer Polytechnic Institute, Troy, NY) ^32^ and were cloned in pRSFDuet-1 plasmid (Novagen, Madison, USA) in multiple cloning site 1 (MCS1). Also, since the UDP-glucose dehydrogenase (UGD) overexpression has been determined to be crucial for glycosaminoglycan production ^39–42^, *Zymomonas mobilis* UGD gene (*Zmugd*) ^42^ was also cloned in the same plasmid in MCS2. The three genes were cloned in pseudo-operon configuration, i.e., the DNA sequence of each gene follows its own lac operator, T7 promoter and RBS, and a single T7 terminator exists in the end of all genes.

**Table 1.**
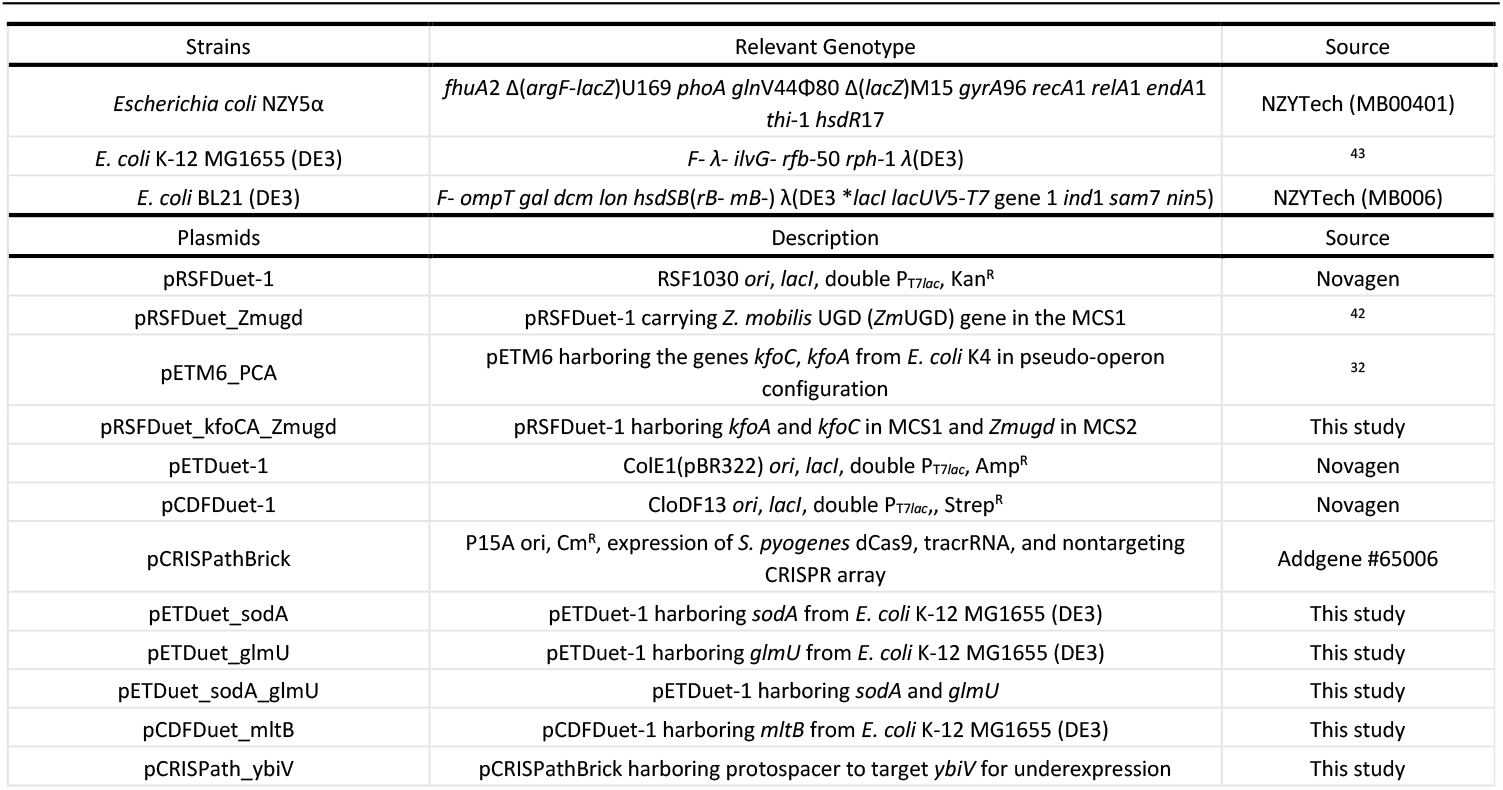
Strains and plasmids used in this study.

Plasmid DNA was extracted using NucleoSpin® Plasmid Miniprep Kit (Macherey-Nagel, Düren, Germany). Genes have been amplified using Phusion High Fidelity DNA Polymerase (Thermo Fisher Scientific, Waltham, USA) and the primers (Metabion, Steinkirchen, Germany; Eurofins, Ebersberg, Germany) are described in Table SI1. Amplified DNA fragments were purified from agarose using NucleoSpin® Gel and PCR Clean-up Kit (Macherey-Nagel). Plasmid DNA and PCR products were quantified using a NanoDrop One instrument (Thermo Fisher Scientific) and were digested with the proper restriction endonucleases (Thermo Fisher Scientific) for 1 h at 37°C and purified using NucleoSpin® Gel and PCR Clean-up Kit. Ligations were performed for 1 h at room temperature using a T4 DNA ligase (Thermo Fisher Scientific). The constructions were transformed by heat shock into *E. coli* NZY5α competent cells (NZYTech, Lisbon, Portugal). Super optimal broth with catabolite repression (SOC; NZYTech) was used for transformants recovery.

All plasmids herein constructed were verified by colony PCR using Dream *Taq* polymerase (Thermo Fisher Scientific) and digestion, and their sequences were further confirmed by sequencing (Eurofins) (Primers in Table SI1). After sequence confirmation, the resulting plasmid with the assembled pathway, pRSFDuet_kfoCA_Zmugd, was transformed in *E. coli* K-12 MG1655 (DE3) and *E. coli* BL21 (DE3) strains for the evaluation of chondroitin production.

### Mutant construction

The evaluated modifications to improve heterologous chondroitin production included membrane-bound lytic murein transglycosylase B (*mltB*), superoxide dismutase (*sodA*) and *N*-acetylglucosamine-1-phosphate uridyltransferase (*glmU*) overexpression, sugar phosphatase (*ybiV*) and lipid II flippase (*murJ*) underexpression, and β-*N*-acetylglucosaminidase (nagZ) deletion. *mltB, sodA* and *glmU* have been amplified from *E. coli* K-12 MG1655 (DE3) genome and cloned using the primers glmU_Fw, glmU_Rv, sodA_Fw, sodA_Rv, mltB_Fw and mltB_Rv (Table SI1) in the same conditions as previously described for chondroitin pathway construction. pETDuet-1 was used as vector to overexpress *glmU* and/or *sodA*, while overexpression of *mltB* was conducted using pCDFDuet-1 as expression vector. pCRISPathBrick plasmid (Table 1) was used to implement the CRISPR interference (CRISPRi) system, by expressing and targeting a dCas9 (dead Cas9) ^44^. The protospacer for *ybiV* and *murJ* targeting was inserted using Golden Gate assembly. For this procedure, annealing of pair of oligos (ybiV_UE_Fw, ybiV_UE_Rv for *ybiV*, Table SI1, and murJ_UE_Fw, murJ_UE_Rv or murJ2_Fw, murJ2_Rv for *murJ*, Table SI2) has been conducted by incubating 100 μmol of each oligo in T4 DNA ligase buffer at 95°C for 5 min and letting the temperature slowly decrease until room temperature was reached. Then the annealed oligo was mixed with the plasmid pCRISPathBrick, *Bsa*I (=*Eco*31I) and T4 DNA ligase. The mixture was incubated through 10 cycles of 5 min at 37°C and 10 min at 22°C, followed by 30 min at 37°C and 15 min at 75°C. All constructed plasmids were verified by colony PCR, digestion and their sequences were further confirmed by sequencing.

In the attempts to delete *nagZ* gene, the protocol for gene deletion using a CRISPR-Cas9 strategy was followed ^45^. The primers used are described in Table SI2.

### Flask fermentations

For chondroitin screening production tests in conical flask fermentations, *E. coli* K-12 MG1655 (DE3) and *E. coli* BL21 (DE3) harboring the heterologous chondroitin production pathway (pRSFDuet_kfoCA_Zmugd) have been used. Also, variations of these strains, with over and/or underexpression of genes, have been evaluated.

Flasks of 250 mL with 50 mL lysogeny broth (LB) Miller (NZYTech) supplemented with proper antibiotics (i.e., depending on the plasmid(s) present in the strain, 50 μg/mL of kanamycin (Fisher), 100 μg/mL of ampicillin (Fisher Bioreagents), 100 μg/mL of spectinomycin (Alfa Aesar) and/or 34 μg/mL chloramphenicol (Alfa Aesar)) have been inoculated with 1% (v/v) of an overnight culture. The cultures were incubated at 37°C and 200 rpm until OD_600nm_ of 0.6-0.8 was reached. At this point, isopropyl β-D-1-thiogalactopyranoside (IPTG, 1 mM; NZYTech) was added to induce heterologous enzyme production and temperature was decreased to 30°C. The cultures were further incubated for 24 h. All assays were conducted in triplicate.

### Bioreactor operation

The engineered *E. coli* K-12 MG1655 (DE3) strains were collected from LB plates supplemented with proper antibiotics and used to inoculate pre-inoculums of 10 mL of LB medium. The cultures were grown at 37°C and 200 rpm for about 22 h. Cells were harvested by centrifugation (4,000 x g, 15 min) and used to inoculate 500 mL conical flasks containing 120 mL of defined medium (per liter: yeast extract 2 g (Labkem, Baldoyle, Ireland), K_2_HPO_4_ 10 g (Panreac, Barcelona, Spain), KH_2_PO_4_ 2 g (Panreac), MgCl_2_ 0.1 g (VWR, Radnor, USA), sodium citrate 0.5 g (Panreac), (NH_4_)_2_SO_4_ 1 g (Labkem), glucose 20 g (Acros Organics, Geel, Belgium) (second pre inoculum) which were then shaken at 200 rpm and 37°C for about 16 h, and used to inoculate the bioreactor with an initial OD_600nm_ of 0.1. Fermentations were performed in a 2-L DASGIP® Parallel Bioreactor System (Eppendorf, Hamburg, Germany). The operating volume for the fermentation was 400 mL of defined medium and proper antibiotics. The temperature set-point was maintained at 37°C, and the pH was automatically controlled at 7 by addition of 2 M NaOH (Sigma, St. Louis, USA). The dissolved oxygen was kept above 30% of saturation by using stirring-speed feedback-control ranging from 250 rpm until 800 rpm and a constant air-flow rate of 0.5 volume air per volume medium per min (12 L/h). After 3 h (OD_600nm_ ∼0.6), 1 mM IPTG was added to the culture and temperature was reduced to 30°C. The fermentation continued until glucose was completely consumed (until ∼54 h). Samples were taken during the fermentation to monitor glucose concentration and OD_600nm_.

For fed-batch fermentations, the batch phase was performed as previously described except that initial glucose concentration was 10 g/L. When glucose concentrations were reduced up to 2-3 g/L, the feeding phase was initiated by feeding a concentrated solution (80 g/L glucose and 18 g/L yeast extract) with a constant feeding rate of 1.5-2.5 mL/h, depending on the glucose consumption rate of the strain. All fermentations were conducted in duplicates.

### Analytic methods

Samples from *E. coli* fermentations were centrifuged to separate cells from supernatant (4,000 x g, 15 min).

The supernatants from the end of flask fermentations were used to quantify extracellular chondroitin while the pellets were further processed to determine intracellular chondroitin.

Pellets (from ∼50 mL of culture) were washed and then resuspended in deionized water (5 mL). The suspended cells were lysed by sonication with a microtip probe linked to Vibra-cell processor (Sonics, Newtown, CT, USA). Keeping the solution on ice during the procedure, short pulses of 3 s ON and 4 s OFF at 30% amplitude were performed until 5 min of active sonication was reached. The resulting lysate was centrifuged (12,000 x g 15 min) to remove insoluble material.

For intracellular chondroitin quantification, the lysates were treated with DNase (New England Biolabs, MA, USA) for 2 h at 37°C and then with proteinase K at final concentration of 2 mg/mL (NZYTech) for 2 h at 56°C. Then, the mixture was boiled for 5 min and centrifuged again (16,000 x g 20 min) to removed insoluble material.

Both extracellular and intracellular samples were precipitated by adding three volumes of cold ethanol and letting the mixture at 4°C overnight. The precipitate was collected by centrifugation at 4,000 x g for 10 min at 4°C. The precipitate was air-dried at room temperature overnight after which it was resuspended in deionized water. Insoluble material was removed by centrifugation (16,000 x g for 20 min). Chondroitin was quantified by uronic acid carbazole assay ^46^ using chondroitin sulfate (Biosynth, Staad, Switzerland) solutions as standards. Samples or standards (125 μL) were mixed with 750 μL sulfuric acid reagent (9.5 g/L sodium tetraborate, Supelco, Bellefonte, USA, dissolved in H_2_SO_4_ > 95%). The mixture was boiled for 20 min. Afterwards, 25 μL of carbazole reagent (1.25 g/L carbazole, Supelco, dissolved in absolute ethanol, Fisher) were added to the boiled samples. The mixture was boiled again for 15 min and cooled down for 15 min. The OD_530nm_ was then read in a microplate reader.

Protein expression in cultured *E. coli* K-12 MG1655 (DE3) carrying pETDuet-1, pETDuet_glmU, pETDuet_sodA, pCDFDuet-1, or pCDFDuet_mltB, was evaluated by sodium dodecyl sulfate polyacrylamide gel electrophoresis (SDS–PAGE) ^42^ (4% stacking gel and 12% running gel). Samples (soluble and insoluble fractions of lysates) were mixed with 2x Laemmli Sample Buffer (65.8 mM Tris–HCl pH 6.8, 2.1% SDS, 26.3% glycerol, 0.01% bromophenol blue and 5% β-mercaptoethanol, from Fisher Scientific, JMGS, Sigma-Aldrich and AppliChem, respectively) and denatured at 95°C for 5 min. The protein marker used was Color Protein Standard—Broad Range (NEB, #77125). After electrophoresis, the gel was stained using Coomassie Blue R-250 (AppliChem) for 15 min and de-stained using distilled water. Samples from bioreactors were collected along the fermentations and analyzed in terms of glucose consumption and cellular growth. Final extracellular chondroitin was quantified using the carbazole method described above. Glucose consumption was monitored along the fermentations through dinitrosalicylic acid (DNS) method ^47^. 3,5-Dinitrosalicylic acid (Acros Organics), sodium potassium tartrate tetrahydrate (Panreac) and NaOH (Sigma) were used to prepare the DNS reagent, which was mixed with same volume of samples, boiled and cooled down by adding deionized water. The OD_540nm_ was then measured. The glucose concentrations were further confirmed by high performance liquid chromatography (HPLC) using a JASCO system associated with a refractive index (RI) detector (RI-2031), and an Aminex HPX-87H column from Bio-Rad, which was kept at 60°C; the mobile phase used was 5 mM H_2_SO_4_ (Fisher) at a flow rate of 0.5 mL/min. All ODs were measured in a 96-well plate spectrophotometric reader Synergy HT (BioTek, Winooski, VT, USA), to establish the growth profile.

## Results and discussion

### Bioinformatic results for mutant prediction

OptFlux was used to identify potential *E. coli* mutants with enhanced capabilities to produce increased quantities of chondroitin. Using this tool, it was not possible to predict the improvement of chondroitin production by combining gene knockouts (*data not shown*). This was likely because the competing pathways that use the intermediates are critical for cell growth. However, performing gene over- and underexpression searches allowed to identify several possible targets. Table 2 displays the genetic modifications identified as potential phenotypes with the highest BPCY for each model.

**Table 2.** Optimization of chondroitin production using OptFlux. The optimization algorithm was run at least four times for each model. The predicted phenotype for the unmodified and modified strains (from the resulting solutions with highest biomass-product coupled yield (BPCY)) are shown. BPCY is calculated by OptFlux by multiplying biomass by product and then dividing by substrate consumed (in all cases being 10 mmol/gDW/h), as predicted by pFBA simulation. Flux variability analysis (FVA) results are shown as minimum and maximum chondroitin obtained through pFBA for fixed biomass. Predicted biomass and chondroitin values are in units of h^-1^ and mmol/gDW/h, respectively.

| Model | BPCY | Genes modified |  | Predicted phenotype (pFBA) |  | FVA |  |
| --- | --- | --- | --- | --- | --- | --- | --- |
|  |  | Underexpression | Overexpression | Biomass | Chondroitin | Min chondroitin | Max chondroitin |
| iB21_1397_c | - | - | - | 0.9756 | 0.0000 | - | - |
| iB21_1397_c | 0.09607 | *cmk* | *glmU* | 0.3671 | 2.6168 | 2.6146 | 2.6345 |
|  | 0.09607 | *pyrH* | *glmM* | 0.3671 | 2.6168 | 2.6146 | 2.6345 |
|  | 0.09607 | *pyrH* | *glmU* | 0.3671 | 2.6168 | 2.6146 | 2.6345 |
|  | 0.09200 | *mltC* | *glmU* | 0.6516 | 1.4119 | 1.4025 | 1.4401 |
|  | 0.09200 | *mltF* | *glmU* | 0.6516 | 1.4119 | 1.4025 | 1.4401 |
|  | 0.09200 | *mltA* | *glmU* | 0.6516 | 1.4119 | 1.4025 | 1.4401 |
|  | 0.09200 | *mltB* | *glmU* | 0.6516 | 1.4119 | 1.4025 | 1.4401 |
|  | 0.09200 | *mltE* | *glmU* | 0.6516 | 1.4119 | 1.4025 | 1.4401 |
|  | 0.09200 | *slt* | *glmU* | 0.6516 | 1.4119 | 1.4025 | 1.4401 |
| iECBD_1354_c | - | - | - | 0.9756 | 0.0000 | - | - |
| iECBD_1354_c | 0.09200 | *mltE* | *glmM* | 0.6516 | 1.4119 | 1.4025 | 1.4401 |
|  | 0.09200 | *slt* | *glmM* | 0.6516 | 1.4119 | 1.4025 | 1.4401 |
|  | 0.09200 | *mltC* | *glmM* | 0.6516 | 1.4119 | 1.4025 | 1.4401 |
|  | 0.09200 | *mltA* | *glmM* | 0.6516 | 1.4119 | 1.4025 | 1.4401 |
|  | 0.09200 | *amiABC, ampG* | *glmU* | 0.6516 | 1.4119 | 1.4025 | 1.4401 |
|  | 0.09200 | *slt* | *glmU* | 0.6516 | 1.4119 | 1.4025 | 1.4401 |
|  | 0.09200 | *mltB* | *glmM* | 0.6516 | 1.4119 | 1.4025 | 1.4401 |
|  | 0.09200 | *mltF* | *glmM* | 0.6516 | 1.4119 | 1.4025 | 1.4401 |
| iEC1356_BI21DE3_c | - | - | - | 0.9767 | 0.000 | - | - |
| iEC1356_BI21DE3_c | 0.09215 | *mltE* | *glmU* | 0.6519 | 1.4135 | 1.4015 | 1.4417 |
|  | 0.09215 | *oppC* | *glmU* | 0.6519 | 1.4135 | 1.3268 | 1.4417 |
|  | 0.09215 | *mltC* | *glmU* | 0.6519 | 1.4135 | 1.4015 | 1.4417 |
|  | 0.09215 | *oppB* | *glmU* | 0.6519 | 1.4135 | 1.3268 | 1.4417 |
|  | 0.09215 | *oppF* | *glmU* | 0.6519 | 1.4135 | 1.3268 | 1.4417 |
|  | 0.09215 | *amiA* | *glmU* | 0.6519 | 1.4135 | 1.4015 | 1.4417 |
|  | 0.09215 | *amiB* | *glmU* | 0.6519 | 1.4135 | 1.4015 | 1.4417 |
|  | 0.09215 | *oppD* | *glmU* | 0.6519 | 1.4135 | 1.3268 | 1.4417 |
|  | 0.09215 | *mltA* | *glmU* | 0.6519 | 1.4135 | 1.4015 | 1.4417 |
|  | 0.09215 | *mltB* | *glmU* | 0.6519 | 1.4135 | 1.4015 | 1.4417 |
|  | 0.09215 | *amiC* | *glmU* | 0.6519 | 1.4135 | 1.4015 | 1.4417 |
| iJO1366_c | - | - | - | 0.9824 | 0.000 | - | - |
| iJO1366_c | 0.09287 | *mltB* | *glmM* | 0.6531 | 1.4219 | 1.4000 | 1.4501 |
|  | 0.09287 | *mltE* | *glmM* | 0.6531 | 1.4219 | 1.4000 | 1.4501 |

Most of the resulting solutions comprised two combined modifications, namely underexpression of one of the genes from cell wall biosynthesis and recycling pathways (such as: lytic transglycosylases MltABCEF and Slt; anhydromuropeptide permease AmpG; oligopeptide permeases *oppBCDF*; or *N*-acetylmuramoyl L-alanine amidases AmiABC) and overexpression of one of the genes responsible for the production of a chondroitin precursor (either glucosamine-1-phosphate *N*-acetyltransferase/ UDP-*N*-acetylglucosamine diphosphorylase GlmU or phosphoglucosamine mutase GlmM). Underexpression of cytidylate kinase *cmk* and UMP kinase *pyrH* genes were also identified, which catalyze reactions from the biosynthesis and salvage of pyrimidine ribonucleotides (Figure 1).

**Figure 1.**
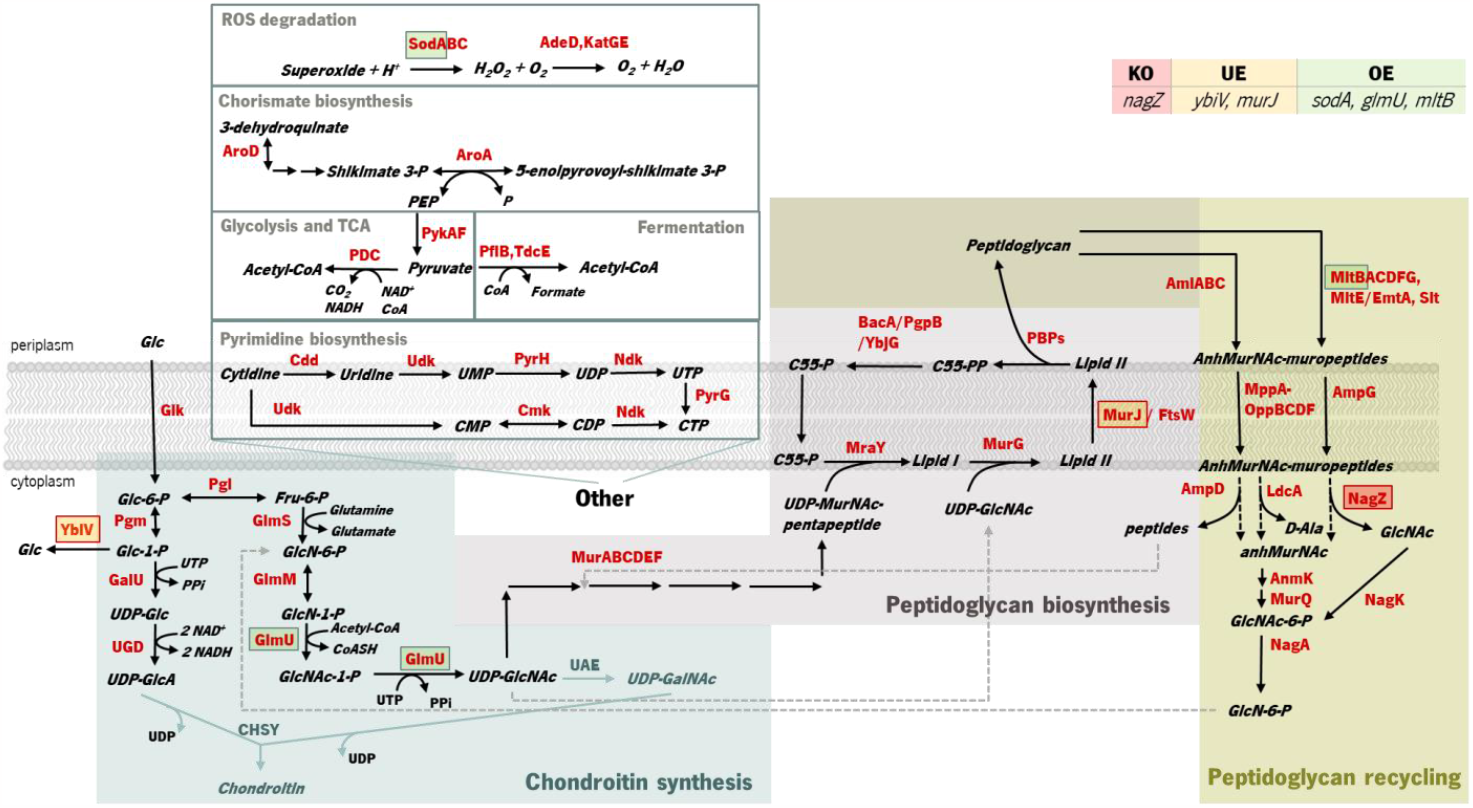
Identified targets for genetic modification to potentially improve chondroitin heterologous production in *Escherichia coli* and their role in bacterial metabolism. The target for knock-out (KO) is marked in a red square, underexpressions (UE) are marked in orange squares, and overexpressions (OE) are marked in green squares. MltB, MurJ and NagZ are involved in cell wall biosynthesis and recycling, while GlmU produces the chondroitin precursor uridine-diphosphate (UDP)-*N*-acetylglucosamine and YbiV dephosphorylates the precursor glucose 1-phosphate. SodA is not directly related with the metabolism of chondroitin synthesis intermediates. Enzyme and compounds abbreviations: AdeD: adenine deaminase; AmiABC and AmpD: *N*-acetylmuramoyl-*L*-alanine amidases; AmpG: anhydromuropeptide permease; AnmK: anhydro-N-acetylmuramic acid kinase; AroA: 3-phosphoshikimate 1-carboxyvinyltransferase; BacA: undecaprenyl pyrophosphate phosphatase; C55: *di-trans,octa-cis*-undecaprenyl; Cdd: cytidine/deoxycytidine deaminase; CHSY: chondroitin synthase; Cmk: cytidylate kinase; FtsW: peptidoglycan glycosyltransferase; GalU: uridine-triphosphate(UTP):glucose-1-phosphate uridylyltransferase; Glk: glucokinase; GlmU: glucosamine-1-phosphate *N*-acetyltransferase/ UDP-*N*-acetylglucosamine diphosphorylase; GlmM: phosphoglucosamine mutase; GlmS: glucosamine-6-phosphate synthase; KatGE: catalases; LdcA: murein tetrapeptide carboxypeptidase; MltABCDFG: membrane bound lytic transglycosylases; MppA-OppBCDF: oligopeptide permeases complex with muropeptide-binding protein; MraY: phospho-*N*-acetylmuramoyl-pentapeptide-transferase; MurG: *N*-acetylglucosaminyl transferase; MurJ: lipid II flippase; MurQ: *N*-acetylmuramic acid 6-phosphate etherase; NagA: *N*-acetylglucosamine-6-phosphate deacetylase; NagK: *N*-acetyl-D-glucosamine kinase; NagZ: β-*N*-acetylglucosaminidase; Ndk: nucleoside diphosphate kinase; PBPs: penicillin-binding proteins; PDC: pyruvate dehydrogenase complex encoded by genes *pdhA, pdhB* and *lpd*; PflB: pyruvate formate-lyase; Pgi: glucose-6-phosphate isomerase; Pgm: phosphoglucomutase; PgpB: phosphatidylglycerophosphatase B; PykAF: pyruvate kinases; PyrG: CTP synthetase; PyrH: UMP kinase; Slt: soluble lytic transglycosylase; SodA: superoxide dismutase; TdcE: 2-ketobutyrate formate-lyase/pyruvate formate-lyase 4; UAE: UDP-*N*-acetylglucosamine 4-epimerase; Udk: uridine/cytidine kinase; UGD: UDP-glucose 6-dehydrogenase; YbjG: undecaprenyl pyrophosphate phosphatase; YbiV: sugar phosphatase.

Overexpression of *glmM* and/or *glmU* are well reported metabolic engineering strategies for improving the production of chondroitin ^48^ or hyaluronic acid ^49^ in *E. coli*.

In Gram-negative bacteria, the cell-wall recycling is initiated in the periplasm by bacterial lytic transglycosylases such as Slt and MltABCEF, that degrade murein (peptidoglycan), the major cell wall component, at the glycosidic bond between *N*-acetylglucosamine (GlcNAc) and *N*-acetylmuramic acid (MurNAc), releasing a distinctive 1,6-anhydro-*N*-acetyl-β-D-muramyl (AnhydroMurNAc) product. The anhydromuropeptide permease AmpG specifically transports the resulting AnhydroMurNAc-containing muropeptides, from the periplasm into the cytoplasm ^50–52^. MurNAc-L-Ala amidases AmiABC, are another class of peptidoglycan degrading enzymes in the periplasm, that cleave the amide bond between MurNAc and the stem peptide in peptidoglycan, releasing murein tri-, tetra-, and pentapeptides into the periplasm from which they diffuse out of the cell, or enter the cytoplasm via the MppA-OppBCDF permease system ^53^. The membrane oligopeptide permeases OppBCDF and the murein tripeptide ABC transporter periplasmic binding protein MppA constitute an ABC transporter which is involved in the recycling of murein tripeptide from either exogenous sources or from amidases action ^53,54^. Once in the cytoplasm, the muropeptides are degraded by amidase AmpD, β-*N*-acetylglucosaminidase NagZ, and LD-Carboxypeptidase LdcA, and their constituent components are used for Lipid II biosynthesis. The Lipid II assembled in the cytoplasm is delivered to the periplasm for the *de novo* synthesis of peptidoglycan ^52^. Consequently, peptidoglycan biosynthesis is the main competing pathway that redirects precursors from chondroitin biosynthesis. Despite these reactions result in the production of the chondroitin precursor glucosamine 6-phosphate (GlcN-6-P) that can also be produced by glucose, they consume the intermediate UDP-acetylglucosamine (UDP-GlcNAc) (Figure 1). Therefore, underexpressing cell wall recycling pathway genes might lead to more available precursors, as predicted by the models.

The pyrimidine nucleotide metabolic process was also identified as a target pathway to be modified to increase chondroitin production. Cmk rephosphorylates cytidylate (CMP) and deoxycytidylate (dCMP). Underexpression of *cmk* (included in an optimization solution using iB21_1397_c model) could lead to improved uridine-5′-triphosphate (UTP) pool as observed in a *cmk* deletion mutant ^55^. This is beneficial for chondroitin production because UTP is a co-factor in chondroitin-precursors biosynthesis steps catalyzed by GalU and GlmU. Uridine monophosphate (UMP) kinase PyrH is also involved in the biosynthesis of pyrimidine nucleotides, specifically catalyzing the phosphorylation of UMP to UDP. In this case, targeting this gene for underexpression, as obtained in an optimization solution using iB21_1397_c model, has not an obvious explanation and no literature was found to support that the underexpression of *pyrH* would improve UTP pool, or in other manner be directly beneficial for chondroitin biosynthesis. However, unexpected targets can have other mechanisms for improving product titers. Targeting genes from purine and pyrimidine biosynthesis pathways for underexpression (including *cmk* and *pyrH*) has been applied for antibody production ^56^ as a growth decoupling strategy to increase product formation. Inhibiting excess biomass formation allows for carbon to be utilized efficiently for product formation instead of growth, resulting in increased product yields and titers ^56^. Interestingly, in Landberg et al. (2020)^56^ study, from the reported 21 selected targets for underexpression, repression of *cmk* consistently resulted in higher product titers. On the other hand, *pyrH* underexpression did not affect production. It is also relevant to mention that the models herein used are stoichiometric models, therefore, they do not comprehend kinetic information that would be valuable for this analysis. The model iB21_1397_c has the UMP kinase reaction set as reversible and as being catalyzed by both *pyrH* and *cmk* encoded enzymes, although it is known that the specific favored reactions are, *in vivo*, the phosphorylation of UMP and phosphorylation of CMP (or dCMP), respectively ^57,58^. Also, *pyrH* is a known essential gene for *E. coli* growth ^59^. Therefore, despite *pyrH* has been identified *in silico* as a target for underexpression, this might not be the best target for improving chondroitin production.

Generally, solutions herein obtained with different models were composed by similar genetic modifications demonstrating the robustness of the modifications associated with the overproduction of chondroitin. Despite allowing for a maximum of 10 modifications, the solution comprised usually only two genetic modifications (the highest number of modifications was five with model iECBD_1354_c, Table 2), thus suggesting that the solutions can be further improved using other approaches.

The FVA analysis shows the range of flux distributions of chondroitin. To evaluate the robustness of a solution, the difference between the minimum and maximum chondroitin production, for a fixed biomass value, should be minimal. The solutions that included oligopeptide permeases *oppBCDF* genes underexpression (iEC1356_Bl21DE3_c, Table 2) resulted in greater difference between minimum and maximum chondroitin production, suggesting that are less robust.

To seek for more robust mutants, a different computational approach was performed using MEWpy ^35^. Using this approach, instead of maximizing one objective function (BPCY), two objective functions were defined, namely the BPCY and WYIELD, which allow to guide the evolutionary algorithm onto more robust solutions. BPCY was calculated by MEWpy by multiplying biomass by product, based on pFBA predictions. WYIELD is the weighed sum of the minimum and maximum product fluxes, constrained to a fixed growth.

Since the model iB21_1397_c achieved the best optimization results (highest BPCY of 0.09607 and highest chondroitin flux of 2.6168 mol/gDW/h), when using OptFlux, this model was selected to use in MEWpy. The same environmental conditions were set, and the evolutionary optimization was run. This procedure resulted in 76 solutions, with 39 identified targets for genetic modification. Type and frequency of each genetic modification throughout all solutions was analyzed in Figure 2. The results from the best solutions obtained using MEWpy are shown in Table 3. Solution 1 and 2 have the highest BPCY score, while solutions 5, 6 and 7 exhibited the highest WYIELD. Phenotype simulation was then performed using OptFlux, with pFBA as simulation method, for each solution and those results are also shown in Table 3.

**Table 3.** Optimization results obtained for iB21_1397_c model using the MEWpy tool in Python, and the corresponding relevant fluxes acc ording to phenotype simulations using parsimonious flux balance analysis (FBA) in OptFlux. BPCY was calculated by multiplying biomass by product and then dividing by substrate consumed, as predicted by pFBA. WYIELD is the weighed sum of the minimum and maximum product fluxes. Flux variability analysis (FVA) results are shown as minimum and maximum chondroitin obtained for fixed biomass. Predicted biomass and chondroitin values are in units of h^-1^ and mmol/gDW/h, respectively.

| Solution | BPCY | WYIELD | Genes modified |  |  | Predicted phenotype (pFBA) |  | FVA |  |
| --- | --- | --- | --- | --- | --- | --- | --- | --- | --- |
|  |  |  | Knock-out | Under expression | Over expression | Biomass | Chondroitin | Min chondroitin | Max chondroitin |
| 1 | 0.08840 | 2.91104 | *nagZ* | *ybiV*, *alsK*, *aroA*, *pflB*, *murJ*, *narU* | *sodA*, *glmU*, *mltB* | 0.3040 | 2.9079 | 2.9034 | 2.9289 |
| 2 | 0.08887 | 2.90887 | *ybiV* | *alsK*, *aroA*, *murJ*, *proA* | *idnT*, *Int*, *glmU*, *gabT*, *malZ* | 0.3056 | 2.9079 | 2.9033 | 2.9220 |
| 3 | 0.04316 | 3.14641 | *nagZ*, *ybiV* | *pflB*, *murJ*, *msbA*, *cmk* | *sodA*, *sapD*, *glmU* | 0.1221 | 2.9150 | 2.9150 | 3.6863 |
| 4 | 0.04316 | 3.14651 | *nagZ* | *ybiV*, *pflB*, *murJ*, *msbA* | *sodA*, *sapD*, *idnT*, *glmU* | 0.1221 | 2.9150 | 2.9150 | 3.6866 |
| 6 | 0.04340 | 2.95361 | *nagZ* | *ybiV*, *alsK*, *aroA*, *murJ*, *pyrE* | *sodA*, *glmU*, *zupT*, *mltB* | 0.1221 | 2.9354 | 2.6382 | 3.6896 |
| Selected genes | 0.08840 | 2.91104 | *nagZ* | *ybiV*, *murJ* | *sodA*, *glmU*, *mltB* | 0.3040 | 2.9079 | 2.9034 | 2.9289 |

**Figure 2.**
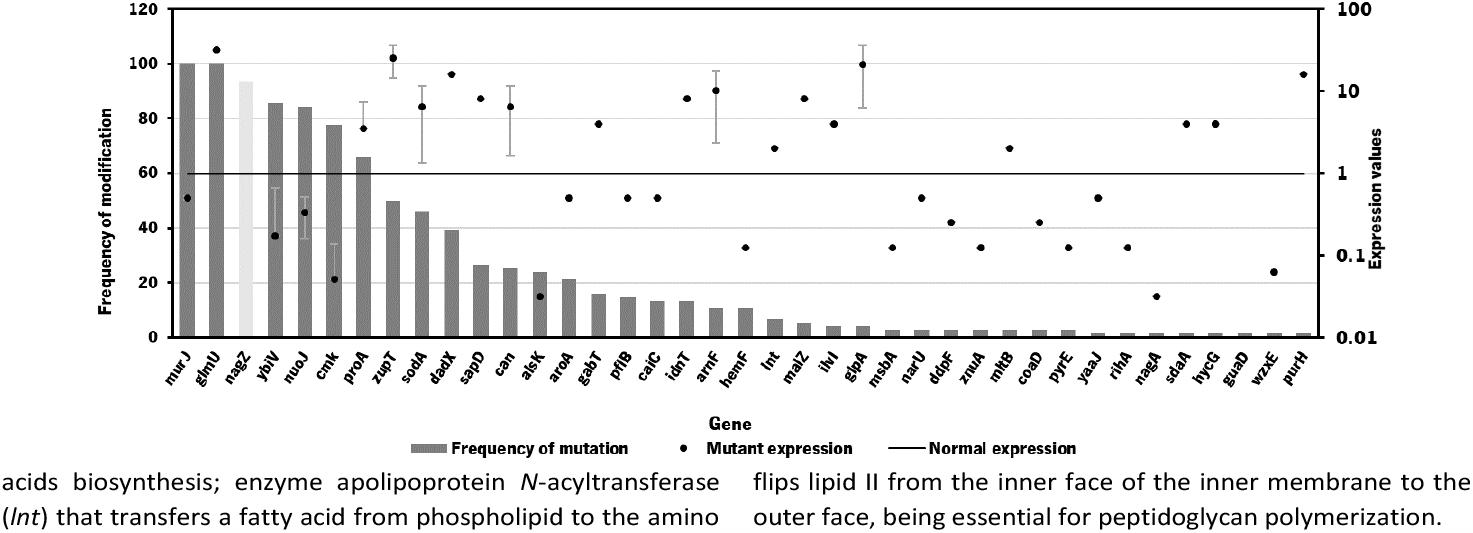
Frequency of genetic modifications in the 76 solutions from strain optimization using MEWpy tool. The mutant expression (in dots) represents the average expression value. Mutant expressions higher than 1 represent overexpression while values of expression lower than 1 represent underexpression. Deletions are represented in light grey bars.

The MEWpy method allowed to identify new mutants with improved chondroitin production, as high as 2.9150 mol/gDW/h, while the highest production obtained from OptFlux solutions was 2.6168 mol/gDW/h (Table 2).

As occurred in the optimization results obtained using OptFlux, solutions from MEWpy included targets from peptidoglycan biosynthesis and recycling pathways (*nagZ, murJ, mltB*) and pyrimidine ribonucleotides salvage pathway (*cmk*). Maltodextrin glucosidase (*malZ*) releases glucose from malto-oligosaccharides as part of the glycogen degradation pathway ^60^, and its suggested overexpression might be to improve glucose and glucose 6-phosphate availability through recycling of those carbon stocks. Many of these new solutions included varied transporters (*narU, znuA, sapD, msbA, idnT*) that were not found to be related with chondroitin production. Other genes that were not found to be related to chondroitin biosynthesis pathway, co-factor production or competing pathways were: D-allose kinase *alsK* that catalyzes the phosphorylation of D-allose to D-allose 6-phosphate; *aroA* that encodes 3-phosphoshikimate 1-carboxyvinyltransferase which is involved in chorismate pathway, leading to aromatic amino acids biosynthesis; enzyme apolipoprotein *N*-acyltransferase (*lnt*) that transfers a fatty acid from phospholipid to the amino terminus of a diacylglycerol prolipoprotein as part of lipoprotein posttranslational modification pathway; *purH* that encodes the bifunctional phosphoribosylaminoimidazolecarboxamide formyltransferase/ inosinic acid cyclohydrolase involved in the *de novo* biosynthesis of purine nucleotides; glutamate-5-semialdehyde dehydrogenase encoded by *proA* that is from the L-proline biosynthesis; 4-aminobutyrate aminotransferase GabT that is involved in 4-aminobutyrate (GABA) degradation. Overexpression of *glmU*, responsible for producing a chondroitin precursor UDP-GlcNAc, was included in most solutions, as occurred in those resulting from OptFlux.

Gene *nagZ* encodes β-*N*-acetylglucosaminidase (NagZ), an enzyme involved in peptidoglycan recycling. NagZ acts specifically by hydrolyzing the β-1,4 glycosidic bond, removing *N*-acetyl-glucosamine (GlcNAc) residues from peptidoglycan fragments that have been excised from the cell wall during growth ^61^. Gene *murJ* encodes for lipid II flippase (MurJ) which flips lipid II from the inner face of the inner membrane to the outer face, being essential for peptidoglycan polymerization.

The gene *mltB*, described above, was identified, in the best solution from MEWpy, as a target for slight overexpression (expression value: 2), contrarily to the solutions obtained in OptFlux, where it has been suggested for underexpression. This might be related with the combination and number of genes being different in the solutions, and *mltB* overexpression might be required to compensate for the negative effect on growth, caused by changes in the expression of several genes. The sugar phosphatase YbiV has been indicated in most solutions for either underexpression or knockout. This is a logical solution as this enzyme redirects metabolic flux from chondroitin production by phosphating glucose-6-phosphate into glucose ^62^ (Figure 1). Pyruvate formate-lyase encoded by *pflB*, which catalyzes non-oxidative cleavage of pyruvate to acetyl-CoA and formate in anaerobically growing cells, was included in some solutions (14%) as underexpressed. The underexpression of *pflB* would decrease the flux through this reaction, leading to an increased availability of pyruvate, and consequently improved glucose-6-phosphate or fructose-6-phosphate levels. In fact, *pflB* deletion has been a common reported target for increased dicarboxylic acid production ^63–65^.

Other common gene identified for overexpression was the superoxide dismutase *sodA*. This gene is expressed in response to oxidative stress and acts by destructing toxic superoxide radicals that are naturally produced during respiratory growth^66^. No direct relationship with chondroitin production improvement was found.

Based on genetic modification frequency, and after confirming that the phenotypes did not vary much from the originally proposed solution (Table 3), the gene selection was narrowed, from genes present in the best solution (Solution 1), for genes with more potential to engineer efficient *E. coli* strains. These selected modifications were: *nagZ* deletion, *ybiV* and *murJ* underexpressions and *sodA, glmU* and *mltB* overexpressions. A schematic representation of the affected pathways with these genetic modifications is presented in Figure 1.

The individual modifications and cumulative modifications of the selected genes were experimentally implemented, to study which gene combinations could benefit chondroitin production the most without compromising *E. coli* growth.

### *In vivo* validation of bioinformatic results

#### Chondroitin production using engineered *E. coli*

The biosynthetic pathway for heterologous production of chondroitin was constructed by cloning the genes *kfoC* and *kfoA* from *E. coli* O5:K4:H4 and *Zmugd* in the plasmid pRSFDuet-1. The amplification products are shown in Figure SI1. The assembled pathway was expressed in both *E. coli* K-12 MG1655 (DE3) and *E. coli* BL21 (DE3) to evaluate chondroitin production (Figure 3).

**Figure 3.**
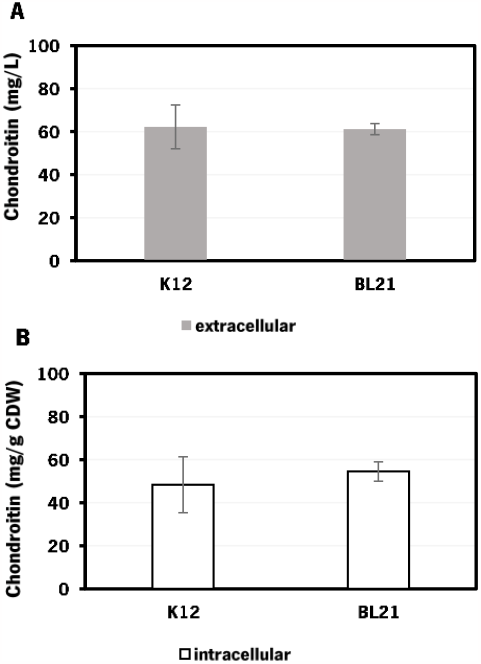
Chondroitin production in *Escherichia coli* engineered strains with chondroitin biosynthetic pathway in flask fermentation: A – extracellular chondroitin in mg/L and B – intracellular chondroitin in mg/g CDW, using K12 – *E. coli* K-12 MG1655 (DE3) and BL21 – *E. coli* BL21 (DE3), both carrying pRSFDuet_kfoCA_Zmugd. Assays were performed in triplicate. CDW – cell dry weight.

*E. coli* K-12 harboring the biosynthetic pathway for chondroitin production was able to produce 62 ± 10 mg/L of extracellular chondroitin and 48 mg/g of cell dry weight (CDW) of intracellular chondroitin. *E. coli* BL21 was able to produce 61 ± 3 mg/L of extracellular chondroitin and 55 mg/g CDW of intracellular chondroitin. These results are higher than the ones obtained in a recent work ^67^ that reported a production of intracellular sulfated chondroitin of 126.64 μg/g CDW and 13.14 μg/g CDW using *E. coli* O5:K4:H4 and *E. coli* K-12 MG1655, respectively, in flasks. These strains harbored a biosynthetic pathway for chondroitin production comparable to the one herein used (*kfoC, kfoA* and *kfoF*, naturally present in *E. coli* O5:K4:H4), but also expressed a chondroitin-4-*O*-sulfotransferase and lacked PAPS reductase *cysH* for the sulfation of chondroitin.

Extracellular chondroitin is more commonly measured, possibly because of its ease of purification and quantification, which makes it of greater interest for biotechnological production. Extracellular chondroitin production using engineered *E. coli* in shake flask cultivation has been reported to achieve concentrations from 0.01076 to 1739 mg/L ^32,67–71^, depending on the host, biosynthetic pathway, chassis optimizations and on culture conditions.

The highest chondroitin titers reported have been obtained using engineered *E. coli* O5:K4:H4, which naturally produces a fructosylated chondroitin, in defined medium, and in batch or fed-batch fermentations in bioreactors. For instance, a three-phase fermentation with pathogenic *E. coli* O5:K4:H4 overexpressing gene from transcription antitermination protein *rfaH* led to 9.2 g/L of chondroitin using glucose as substrate, and in a larger scale, the same strain produced 9 g/L using glycerol as substrate ^68^. To avoid the risks of using pathogenic bacteria, there have been efforts to construct alternative hosts to be efficient for chondroitin production using metabolic engineering strategies. A *Bacillus subtilis* 168, engineered with *kfoC, kfoA* expression and *tuaD* up-regulation, growing on sucrose, has achieved 5.22 g/L of chondroitin in fed-batch fermentation ^39^.

As the culture medium herein used for these initial screening tests in flasks was LB, the optimal production of chondroitin could be further optimized using different culture conditions. Based on work with highest reported chondroitin production ^68^, a defined medium was used for further fermentations in bioreactor.

As the difference in chondroitin production between the two hosts was not significant (Figure 3), the *E. coli* K-12 MG1655 (DE3) was selected for further engineering strategies.

#### Construction of engineered *E. coli* strains based in bioinformatics optimization

Based on the bioinformatics results, the selected modifications to be implemented towards an enhancement of chondroitin production were the overexpression of *sodA, glmU* and *mltB*; the deletion of *nagZ*; and the underexpression of *ybiV* and *murJ*. The genes *sodA, glmU* and *mltB* for overexpressions were amplified (Figure SI2A) and cloned in pETDuet-1 or pCDFDuet-1. The plasmid with lower copy number (pCDFDuet-1) was used for *mltB* since, from the overexpressions predicted, it was the one with lower expression value (Figure 2). Their expression was confirmed by SDS-PAGE (Figure SI2B).

The *nagZ* knockout was attempted multiple times in *E. coli* K-12 MG1655 (and afterwards in *E. coli* BL21) using a CRISPR-Cas9 strategy ^45^ (schematized on Figure SI3), but it was unsuccessful. The primers used for this strategy are described in Table SI2. Although this gene is reported as non-essential for *E. coli* growth and has been previously deleted ^72^ using a different recombination-based strategy ^73^, it has a described role in cell wall biosynthesis as was previously mentioned. Therefore, it is possible that a strain lacking *nagZ* could be more susceptible to the antibiotics used as selective markers in the attempted CRISPR-Cas9 strategy (spectinomycin and chloramphenicol). A deletion of this gene could affect the cell wall integrity or permeability, that is maintained by the coordinated and regulated action of enzymes involved in the peptidoglycan synthesis and recycling ^74^. In fact, the improved sensibility to β-lactam antibiotics in Gram-negative strains lacking *nagZ* compared to wild-type strains has been widely reported ^61,74,75^. If the cells are more susceptible to the antibiotics used in the selection medium, then it could affect the growth and survival of the cells during the gene editing process. The selective pressure applied by the antibiotics could be too strong, resulting in a decrease in the number of cells that survive and grow on the selection medium. One possible solution to this issue could be to use lower concentrations of the antibiotics to reduce the effect on growth of cells lacking *nagZ*. The integration of the chondroitin pathway genes in the genome without maintenance of antibiotic resistance markers can also be an efficient strategy to reduce the toxic effect of antibiotics. Regarding genes underexpressions, a CRISPRi system ^44^ was designed and constructed. In this strategy, a modified version of the caspase 9 protein commonly called as dead Cas9 (dCas9), which does not have the nuclease activity but maintains sequence-specific double stranded DNA-binding capability, is expressed to target the gene, ultimately repressing its expression. The CRISPRi system for *ybiV* underexpression was successfully constructed and evaluated. However, underexpression of *murJ* was not evaluated because construction of targeting protospacer has failed. A schematic representation of the strategy used can be found on Figure SI3. The primers used in the attempts of constructing the protospacer for *murJ* underexpression are described in Table SI2.

The individual and cumulative genetic modifications (*sodA, glmU*, and *mltB* overexpressions and/or *ybiV* underexpression) were further evaluated *in vivo*.

#### Chondroitin production in shake flasks using *E. coli* engineering strains based in bioinformatic optimizations

The solutions obtained though bioinformatics were further validated *in vivo*. Each modification was individually evaluated in the engineered *E. coli* strain harboring the chondroitin pathway already expressing 3 heterologous genes (*kfoC, kfoA* and *Zmugd*), through shake flasks fermentations (Figure 4) to seek for mutants with improved chondroitin production carrying as few modifications as possible.

**Figure 4.**
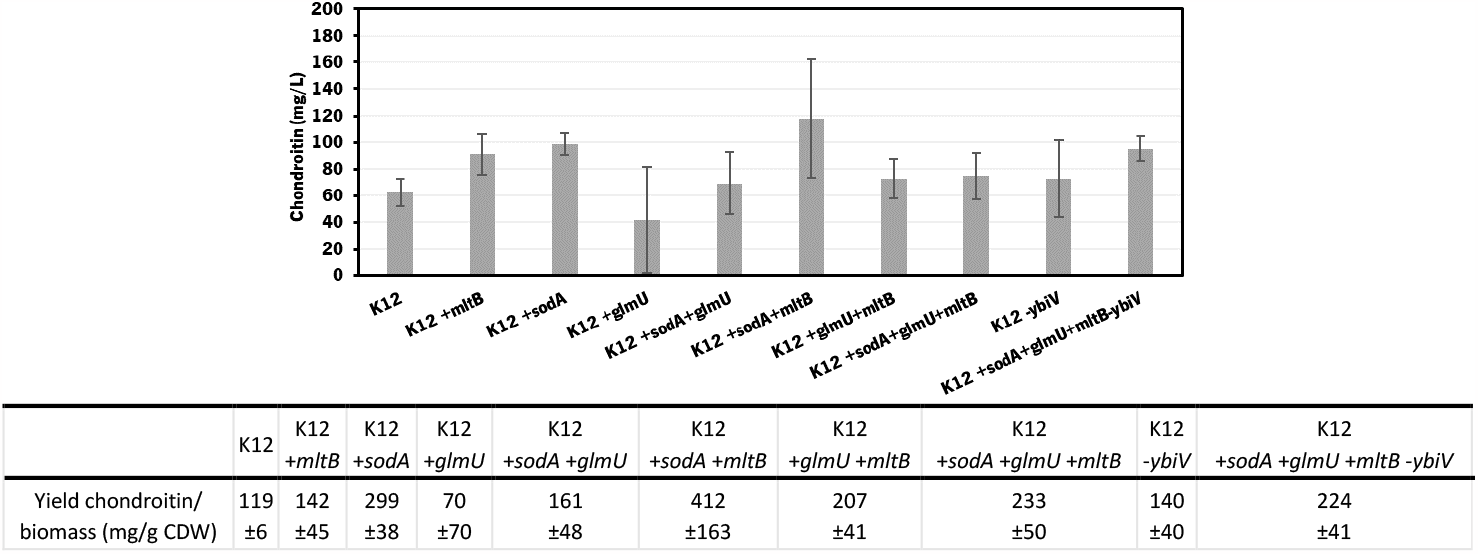
Chondroitin production in *E. coli* K-12 MG1655 harboring the chondroitin biosynthetic pathway (K12, genes *kfoC, kfoA* and *Zmugd*) with additional modifications: *sodA, glmU* and *mltB* overexpressions and/or *ybiV* underexpression. The table shows a comparison of chondroitin yield related to cell dry weight (CDW), obtained for the different mutants.

In these screening shake flask experiments, *E. coli* K-12 mutants were able to produce extracellular chondroitin from 42 to 118 mg/L. Even though the differences in chondroitin production were not significant (*p*-value > 0.05), mutants overexpressing *sodA* or *mltB*, or the one overexpressing both these two genes, seemed to be the most promising ones. As the cumulative effect of both overexpressions was not significantly better than the individual mutations, the two mutants containing individually overexpressed *sodA* or *mltB* were selected for further scale-up studies, in a more suitable culture medium for chondroitin production.

#### Chondroitin production in bioreactor (Batch experiments)

The two selected mutant strains were cultured at bioreactor scale, in batch mode, starting with 20 g/L of glucose. The performance of both strains was further compared to the control (*E. coli* K-12 harboring pRSFDuet_kfoCA_Zmugd). The strain growth and the glucose consumption were monitored (Figure 5).

**Figure 5.**
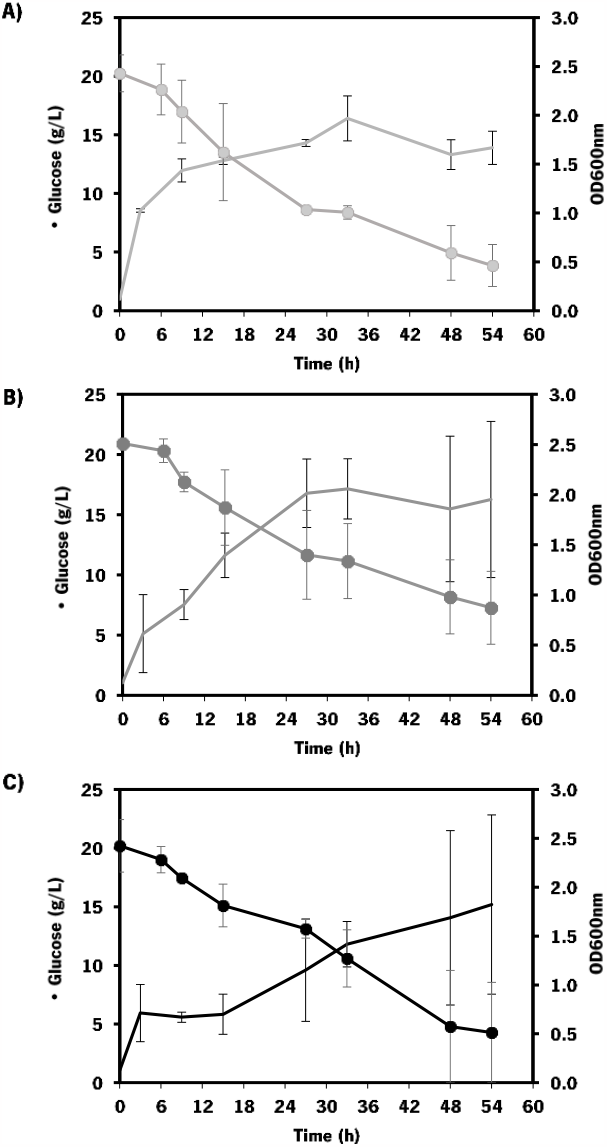
Growth and substrate consumptions curves of *E. coli* K-12 MG1655 DE3 mutants: **A)** carrying pRSFDuet_kfoCA_Zmugd (control - only containing chondroitin biosynthetic pathway); **B)** carrying pRSFDuet_kfoCA_Zmugd and pETDuet_sodA; **C**) carrying pRSFDuet_kfoCA_Zmugd and pCDFDuet_mltB. Data represents average values and standard deviation of two independent experiments. Batch assays starting with 20 g/L glucose. Dots (•) indicate glucose concentration and lines (-) optical density at 600 nm (OD600nm).

The mutants with the selected overexpressions showed more variability between assays but the growth curve was similar to the control which indicates that the growth was not significantly affected by the additionally introduced modifications.

The chondroitin production in the end of fermentation (54 h) was evaluated for each engineered strain (Figure 6).

**Figure 6.**
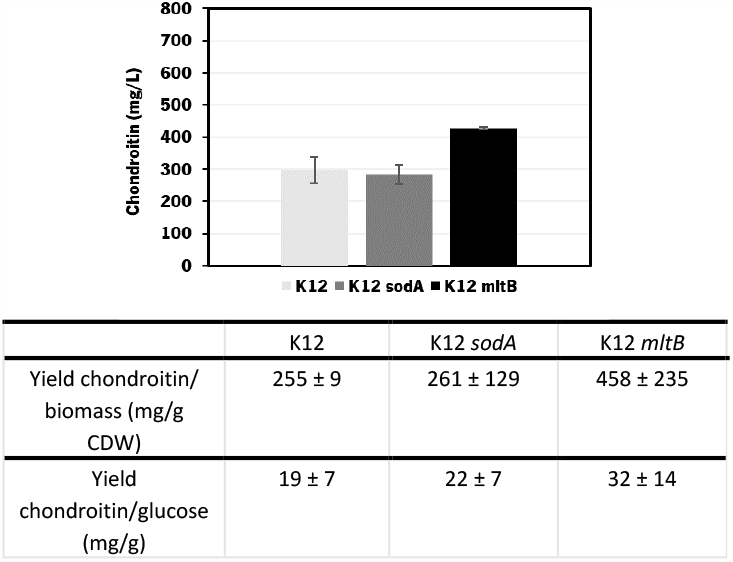
Chondroitin production from cultured *E. coli* K-12 MG1655 (DE3) harboring pRSFDuet_kfoCA_Zmugd (K12) and its counterparts with additional *sodA* or *mltB* overexpressions, in bioreactors operated in batch mode. The table shows a comparison of yield of chondroitin related to cell dry weight (CDW) and yield of chondroitin related to glucose, obtained for the different mutants.

The results obtained showed that *E. coli* K-12 MG1655 engineered with chondroitin production pathway and *mltB* overexpression performed better in terms of extracellular chondroitin production, achieving a concentration of 427 ± 4 mg/L in 54 h. Regarding the yields per biomass or substrate consumption, both mutants expressing *mltB* or *sodA* showed better results compared to the control host. Although the differences between the control and *sodA* mutant are not significant (*p-*value > 0.05), *mltB*-overexpressing strain had improved yields of 1.7-fold, with statistically significant differences (*p*-value < 0.05) compared to the control host or to the *sodA* expressing mutant.

### Chondroitin production in bioreactor (Fed-batch experiments)

The most promising mutant (*E. coli* K-12 MG1655 overexpressing *kfoC, kfoA, Zmugd* and *mltB*) was further cultivated under fed-batch conditions and compared to the control (strain without *mltB* overexpression). The growth and substrate consumption are described on Figure 7.

**Figure 7.**
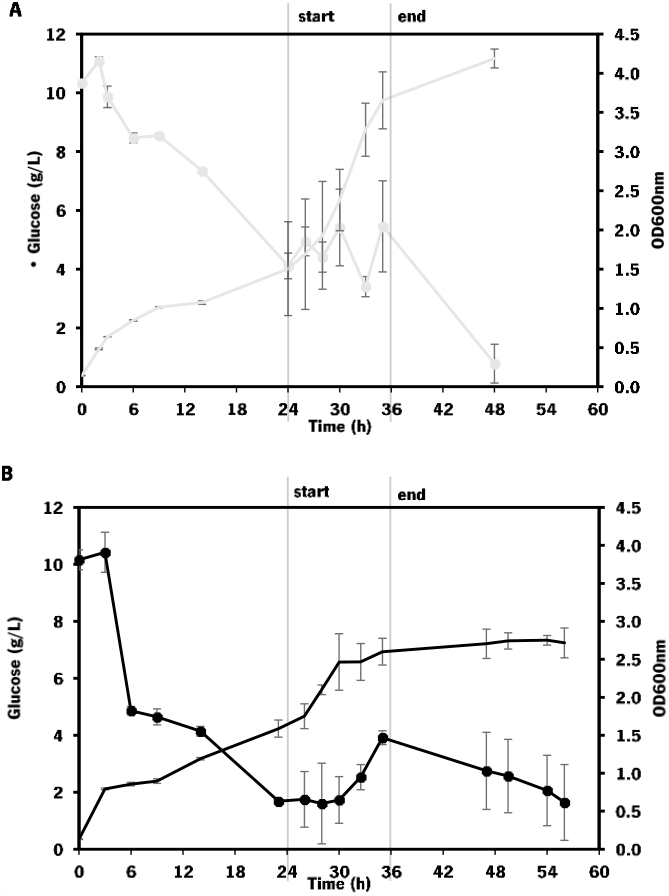
Growth and substrate consumptions curves of: **A:** *E. coli* K-12 MG1655 (DE3) control (only containing chondroitin biosynthetic pathway - pRSFDuet_kfoCA_Zmugd); **B:** *E. coli* K-12 MG1655 (DE3) carrying pRSFDuet_kfoCA_Zmugd and pCDFDuet_*mltB*. Data represents average values and standard deviation of two independent experiments. Fed-Batch assays starting with 10 g/L glucose. The feeding was started after approximately 24 h until 36 h after inoculum.

As expected, the growth of engineered *E. coli* K-12 in bioreactor was greatly improved by changing to fed-batch mode, comparing to the fermentations in batch operation mode. However, this effect was more evident on the control strain (which showed a 2.1-fold increase in growth) than on the one overexpressing *mltB* (that exhibited only a 1.2-fold increase in growth).

The chondroitin production in the end of fermentation was also evaluated (Figure 8).

**Figure 8.**
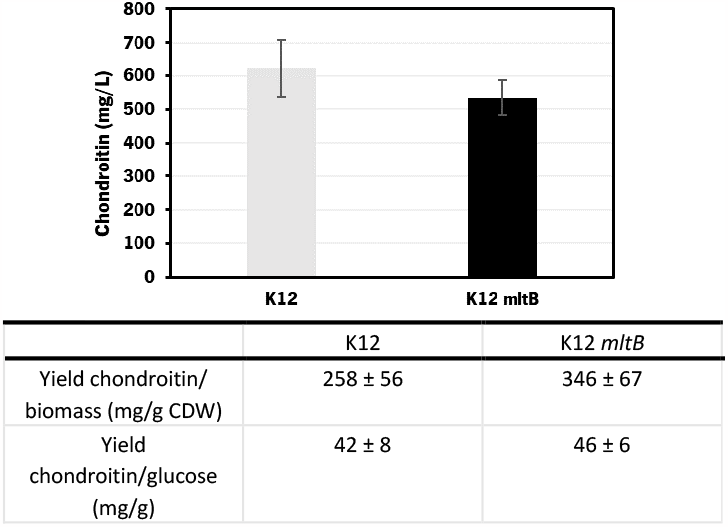
Chondroitin production from cultured *E. coli* K-12 MG1655 harboring pRSFDuet_kfoCA_Zmugd (K12) and its counterpart with additional *mltB* overexpression, in bioreactors operated in fed-batch mode. The table shows a comparison of yield of chondroitin related to cell dry weight (CDW) and yield of chondroitin related to glucose, obtained for the different mutants.

Using the *E. coli* K-12 MG1655 engineered strains lacking or containing *mltB* overexpression, chondroitin concentration at the end of fed-bath fermentation achieved 621 ± 85 mg/L and 535 ± 52 mg/L, respectively. Despite the chondroitin production being lower in the mutant strain overexpressing the *mltB* gene, the yields on biomass and on glucose were higher (1.3-fold and 1.1-fold, respectively) compared to the parent mutant. This is justified with the lower growth of the *mltB*-expressing strain previously discussed (Figure 7).

Chondroitin production has benefited from changing the operation mode to the fed-batch for both strains, resulting in 2.1 and 1.3 -fold increase in chondroitin titers, for control and *mltB* strains, respectively. This is in accordance with other works in the literature that showed that fed-batch fermentations achieved the higher titers comparing to batch or shake flasks fermentations ^32,68,69,76^. Although yields on biomass for both strains were lower than the ones obtained with batch, the yields on glucose were 2.2 and 1.4 times improved, for control and *mltB* strain, respectively. It is common for the fed-batch fermentations to result in much higher cell density, which occurred in this work, and if the growth is not accompanied by product formation at the same range, the yields in biomass are naturally lower. Nevertheless, when comparing both yields on biomass and on substrate, it is evident that *mltB*-overexpressing strain was consistently the best performer.

Although *mltB* overexpression was not a logical modification to improve chondroitin production, it has been predicted in some solutions from computational optimization, including the one with highest BPCY (Solution 1, Table 3). In the batch experiments at bioreactor scale, the *mltB*-overexpressing mutant was indeed the best performing strain in terms of chondroitin production and its growth capability was similar to the strain containing only the chondroitin pathway, without further modifications. However, in fed-batch, the mutant with *mltB* up-regulation did not grow as much as the control. This can be due to the fact that the *E. coli* host used was already expressing three heterologous genes for producing chondroitin, and further *mltB* overexpression might have constrained the bacterium growth as a result of an increase in the metabolic burden, which became more significant when higher cell densities were achieved (fed-batch). We believe that the chondroitin production by these strains could be further improved and become more reproducible with gene integration into *E. coli* genome, rather than being overexpressed using plasmids.

The strain overexpressing *mltB* presented the best chondroitin yields in bioreactors when operated in both batch and fed-batch modes, which suggests that a slight improvement in the peptidoglycan recycling can redirect the metabolic flux towards the chondroitin precursors production.

## Conclusions

In the current study, genome-scale metabolic models’ optimizations were used to identify genes for under- and overexpression, which allowed the selection of possible targets to improve chondroitin production. The suggested promising mutants were further validated *in vivo* by constructing the *E. coli* mutant strains containing the chondroitin heterologous pathway and the additionally selected modifications. In flask 13 fermentation, *E. coli* harboring the biosynthetic pathway was able to produce 62 mg/L of chondroitin. The evaluated mutants 14 with additional modifications on this engineered strain resulted 15 in chondroitin titers from 42 to 118 mg/L. In bioreactor, batch fermentations led to an enhanced chondroitin production, with 16 the highest titer achieved by *E. coli* K-12 overexpressing *mltB* (427 mg/L in 54 h). Further fed-batch assays resulted in an improvement up to 535 mg/L of chondroitin production. This 17 study highlights new possible metabolic engineering targets to improve chondroitin production which ultimately can 18 contribute to advancing the biotechnological production of this most sought glycosaminoglycan.

## Conflicts of interest

There are no conflicts to declare. 21

## Acknowledgements

This study was supported by the Portuguese Foundation for Science and Technology (FCT) under the scope of the strategic funding of UIDB/04469/2020 unit. The authors acknowledge FCT for funding the doctoral grant SFRH/BD/132998/2017 and further extension COVID/BD/152454/2022 to Márcia R. Couto.

## Notes

### Competing Interest Statement

The authors have declared no competing interest.

