## Supplementary file for "Optimization of chondroitin production in *E. coli* using genome scale models"

Supplementary Information

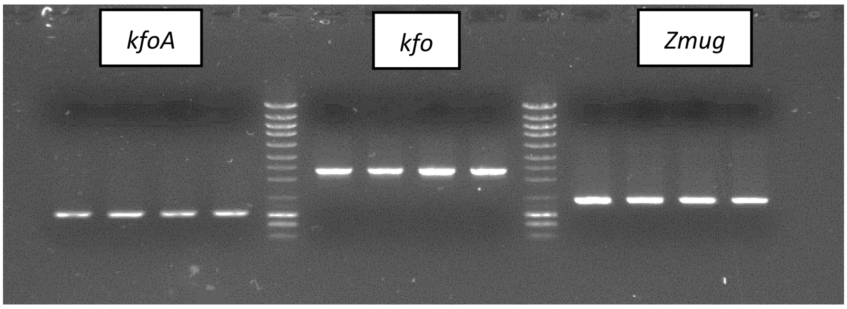


**Figure SI1.** Agarose gel 0.7% showing polymerase chain reaction (PCR) amplification results of genes for chondroitin biosynthetic pathway construction. Ladder: NZYDNA Ladder III, NZYTech.

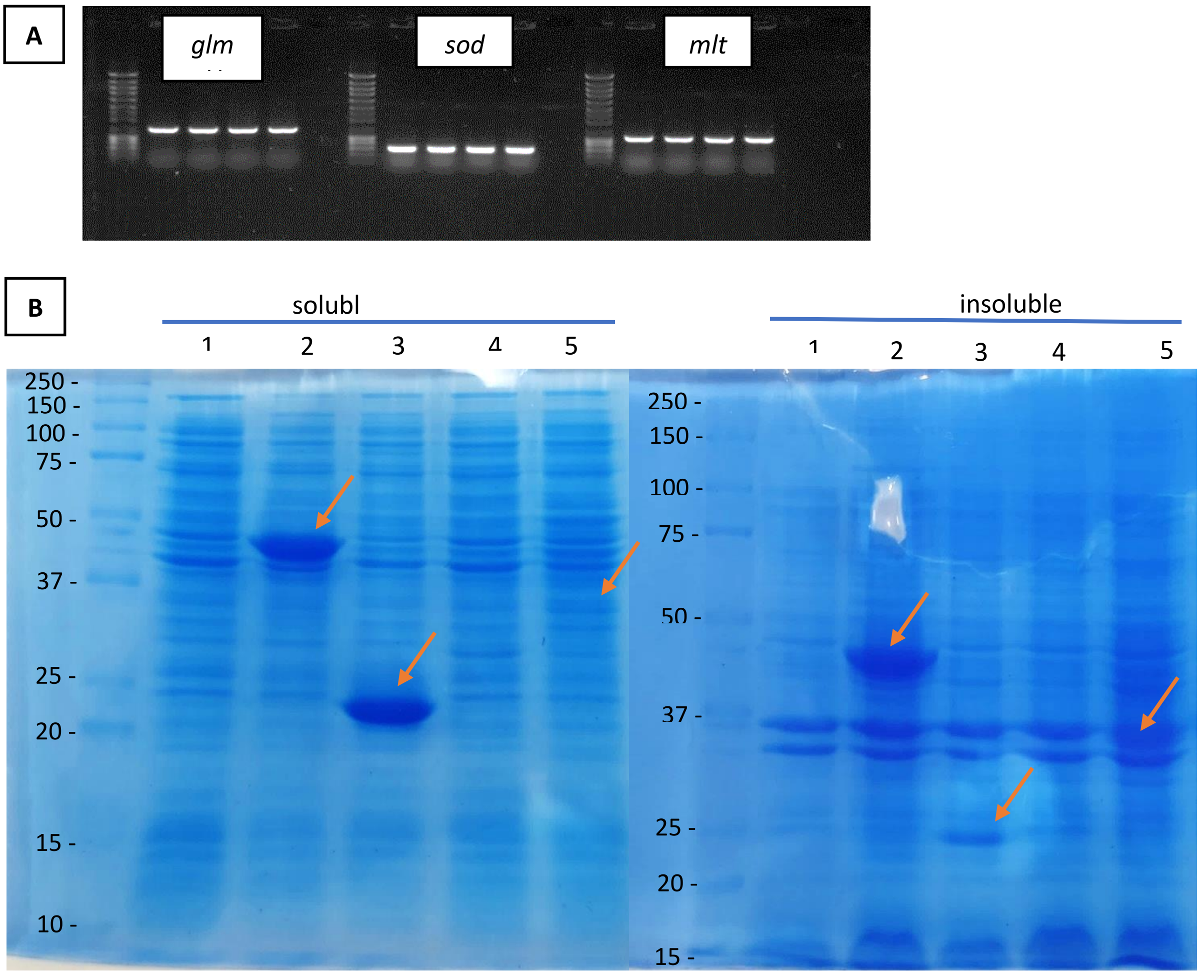


**Figure SI2**. Cloning and expression of identified targets for chondroitin production enhancement: *glmU*, glucosamine-1-phosphate acetyltransferase/*N*-acetylglucosamine-1-phosphate uridyltransferase; *mltB*, membrane-bound lytic murein transglycosylase; *sodA*, superoxide dismutase. **A.** Agarose gel 0.7% showing PCR result of amplification of genes *glmU*, *sodA* and *mltB* from *E. coli* K-12 MG1655 (DE3) genome. **B.** SDS-PAGE gel showing overexpression of genes *glmU*, *sodA* and *mltB* in *E. coli* K-12 MG1655 (DE3). 1 – pETDuet-1, 2 - pETDuet_glmU, 3 – pETDuet_sodA, 4 – pCDFDuet-1, 5 – pCDFDuet_mltB. The predicted sizes were: GlmU 49.2 kDa; SodA 24.91 kDa; MltB 41.2 kDa (Slt35 36 kDa).

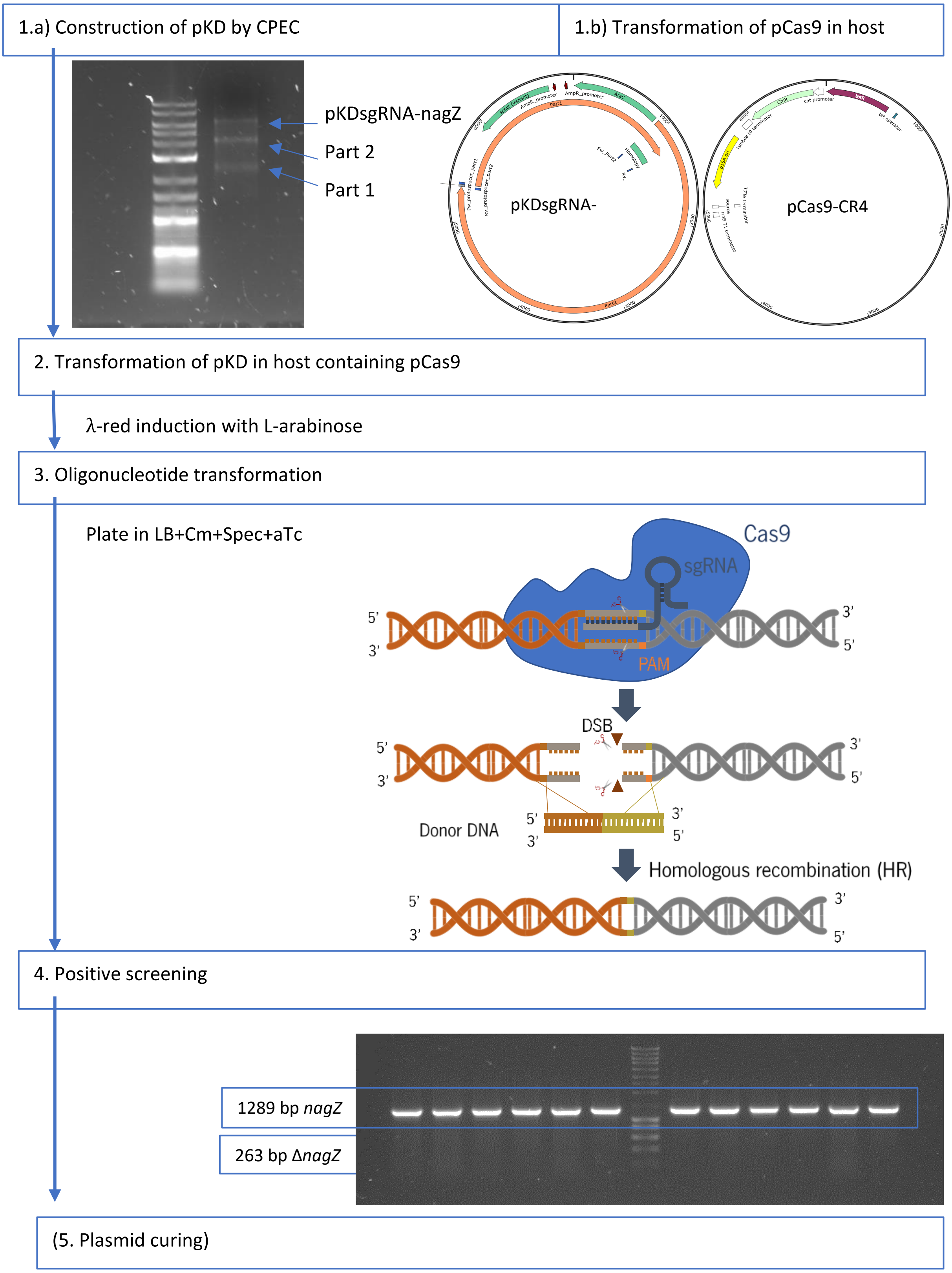


**Figure SI3.** Attempted CRISPR-Cas9 strategy for *nagZ* deletion. This methodology consists in the cleavage of double stranded DNA by Cas9 and λ-Red recombinase system-facilitated genomic integration of donor DNA. The single-guide RNA (sgRNA) encoding plasmid pKDsgRNA-nagZ to target the *nagZ* has been constructed. The plasmids for the expression of Cas9 and of the sgRNA were transformed. Then, the transformation of the oligonucleotide (donor DNA) that should induce the gene deletion was performed. During the subsequent screenings, no positive colonies were found despite several attempts of this procedure.

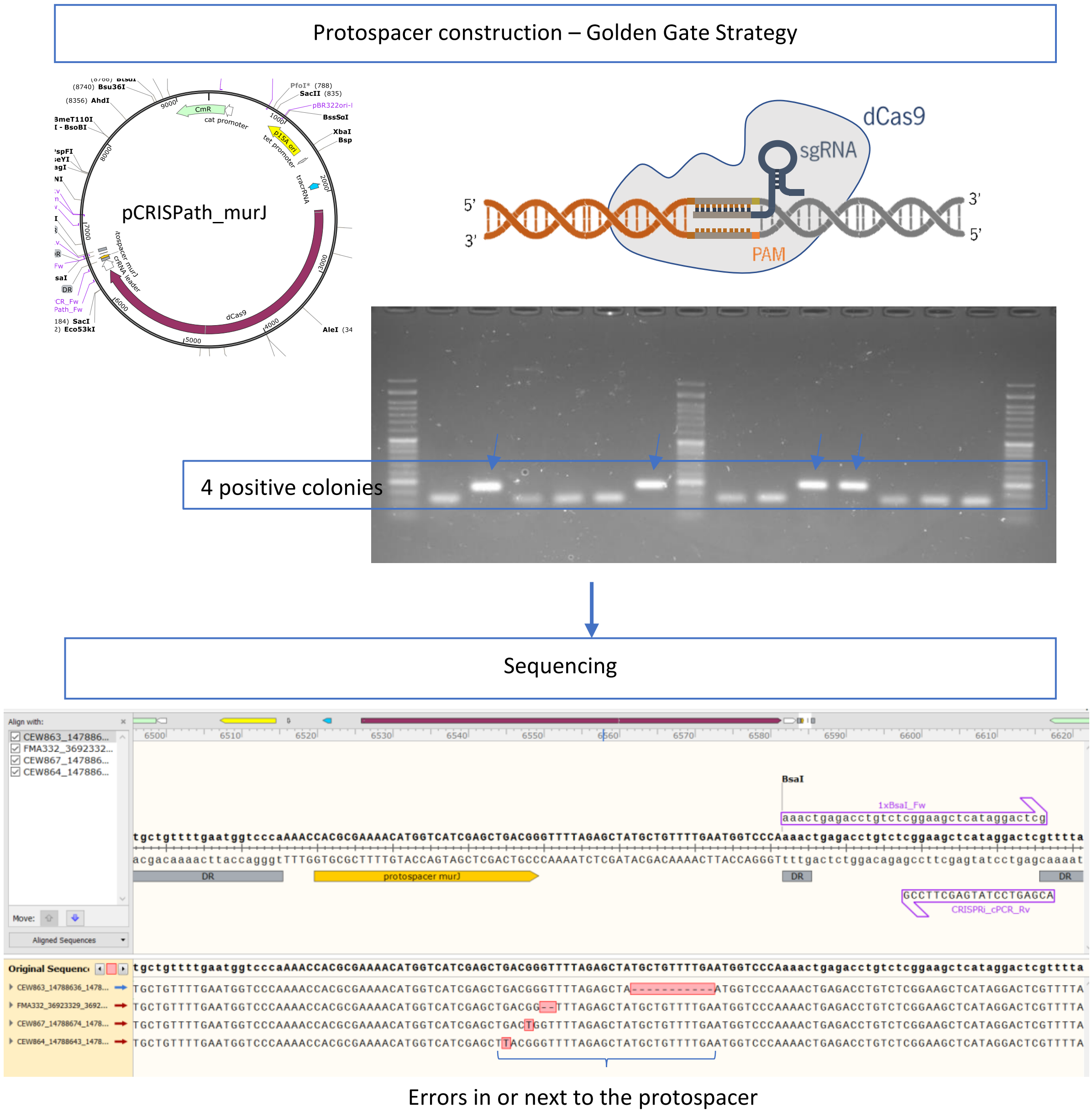


**Figure SI4.** Attempted CRISPR interference (CRISPRi) strategy to underexpress *murJ*. In this strategy, a modified version of the caspase 9 protein, commonly referred as dead Cas9 (dCas9), is expressed to target the *murJ* gene, ultimately repressing its expression. The dCas9 variant lacks nuclease activity but maintains the capability to specifically bind to double stranded DNA sequences. The cloning of the protospacer in the pCRISPathBrick was performed by Golden Gate strategy. The resulting pCRISPath_murJ revealed that the constructed plasmids constantly exhibited errors in or next to the protospacer.
